# Different modality-specific mechanisms mediate perceptual history effects in vision and audition

**DOI:** 10.1101/2022.08.07.503081

**Authors:** Irene Togoli, Michele Fornaciai, Domenica Bueti

## Abstract

Perceptual history plays an important role in sensory processing and decision making, shaping how we perceive and judge external objects and events. Indeed, past stimuli can bias what we are currently seeing in an attractive fashion, making a current stimulus to appear more similar to its preceding one than it actually is. Such attractive effects across successive stimuli appear to be ubiquitous, affecting almost every aspect of perception – from very basic visual attributes (i.e., orientation) to more complex features (i.e., face identity) – suggesting that they may reflect a fundamental principle of brain processing. However, it is unclear whether the ubiquitous nature of these effects is due to an underlying centralised mechanism mediating all of them, or by the existence of separate mechanisms implemented independently in different perceptual pathways. Here we address this question by assessing the behavioural and neural signature of perceptual history in audition and vision, in the context of time perception. Our results first show a double dissociation between the two modalities, whereby the behavioural effect of perceptual history shows opposite patterns of selectivity for the features and position of the stimuli. Electroencephalography results further support a difference between audition and vision, demonstrating that the signature of perceptual history unfolds according to different dynamics in the two modalities and show different relations with the behavioural effect. Overall, our results suggest that the effect of perceptual history may be mediated by different and at least partially independent mechanisms based on the same computational principle, implemented in different sensory pathways.

**SIGNIFICANCE STATEMENT:** The recent history of stimulation, or perceptual history, plays a fundamental role in perception, shaping what we see according to what we saw in the past. The brain mechanisms mediating the integration of past and present perceptual information are however still unclear. In this study we asked whether perceptual history operates via a centralized mechanism shared across sensory modalities, or via distinct modality-specific mechanisms. Our findings show a double dissociation in attractive perceptual history effects across vision and audition, while EEG data show neural signatures of perceptual history with distinct dynamics and properties. Overall, we thus demonstrate that perceptual history affects sensory processing starting from the earliest level of processing, within distinct modality-specific sensory pathways.

## INTRODUCTION

What we perceive in any given moment is not solely determined by the physical information reaching our sensory organs, but it is strongly influenced by the temporal context in which such information is embedded in. Indeed, the recent history of stimulation, or *perceptual history* (i.e., information from the stimuli processed in the recent past) has been shown to shape how we perceive and judge stimuli in the external environment. One classic example of how a past stimulus affects what we perceive in the present is the process of perceptual adaptation, whereby a long exposure to a stimulus causes the perception of the subsequent one to be strongly repulsed away from it (e.g., see Kohn, 2007 for a review). Differently from adaptation, briefer exposures to sensory stimuli can induce attractive biases across successive stimuli, effectively making a stimulus to appear more similar to its preceding one than it actually is (e.g., (J. Fischer & Whitney, 2014)). This attractive influence, often named *serial dependence*, has been shown to be nearly ubiquitous in vision, spanning from basic perceptual dimensions such as orientation (C. Fischer et al., 2020; J. Fischer & Whitney, 2014; Pascucci et al., 2019), duration (Togoli et al., 2021) or numerosity (Corbett et al., 2011; Fornaciai & Park, 2018b), to more complex attributes such as face identity (Liberman et al., 2014). Although most of the studies concerning serial dependence effects focused on vision, similar biases have been also demonstrated in audition (e.g., Motala et al., 2020), although evidence in this modality is much more limited compared to visual perception.

While the increasing number of studies concerning perceptual history has progressively refined our understanding of its properties, pinpointing the exact nature of this effect and its neural mechanisms has proven to be a difficult task. Several ideas have been advanced so far to explain the underlying mechanisms of this phenomenon, proposing that either perceptual history effects occur at the sensory or perceptual level (i.e., thus genuinely biasing what we perceive; Cicchini et al., 2017; Manassi et al., 2018), or occur at post-perceptual processing stages (i.e., thus biasing either memory or decision making; Fritsche et al., 2017; Pascucci et al., 2019). For instance, perceptual history has been interpreted as a signature of a perceptual mechanism supporting visual stability and continuity (e.g., Fischer & Whitney, 2014; Manassi et al., 2018; Murai & Whitney, 2021). Alternatively, the attractive effect has been linked to a higher-level mechanism based on decision-making and memory, depending on the judgment of past stimuli rather than perceptual history per se (e.g., Bliss et al., 2017; Pascucci et al., 2019). Both these frameworks are supported by different findings suggesting that either the attractive effect is perceptual (Cicchini et al., 2017; Fornaciai & Park, 2018a, 2020a) or decisional in nature (Pascucci et al., 2019; Wehrman et al., 2020). Overall, there is evidence supporting both the perceptual and cognitive accounts. Indeed, the behavioural attractive bias shows many properties of high-level processing (i.e., generalization across very different stimuli, influence of confidence; Fornaciai & Park, 2019, 2022; Samaha et al., 2019). Conversely, other studied showed that the effect even precedes other perceptual biases (Cicchini et al., 2021), and electroencephalography studies showed a neural signature of perceptual history at very early latencies after the onset of a stimulus (i.e., 50-100 ms; Fornaciai et al., 2020; Fornaciai & Park, 2018a, 2020). The neural mechanisms governing the implementation of perceptual history and the serial dependence effect thus remain mostly unclear.

An important feature of perceptual history that may help understanding its mechanism is its widespread nature across several perceptual domains and sensory modalities. Considering this feature, the question thus is: does the implementation of perceptual history and the related serial dependence bias involve a unique, generalised mechanism shared across dimensions and modalities, or does it involve a series of distinct mechanisms implemented independently in different sensory pathways?

Answering this question would also allow to better discern the nature of perceptual history effects and the underlying neural mechanisms. Indeed, while the idea of a purely high-level, decisional mechanism would predict an underlying centralised mechanism, a more perceptual mechanism mediating the implementation of perceptual history would instead predict the involvement of low-level sensory-specific processing pathways. To address these possibilities, we tested and compared the attractive serial dependence effect and the neural (electrophysiological) signature of perceptual history in time perception, across audition and vision.

In Exp. 1, we investigated the properties of the behavioural attractive effect in audition and vision, focusing on two important aspects of the effect: the selectivity for the features of successive stimuli (i.e., whether the effect generalises across stimuli with different contextual features; feature selectivity condition) and the selectivity for the spatial position of the stimuli (i.e., whether the effect works across different spatial locations; spatial selectivity condition). In Exp. 2, we replicated the same psychophysical paradigm, but with the addition of electroencephalography (EEG) to disentangle the neural signature of perceptual history in the two modalities.

## RESULTS

### Experiment 1

To address the mechanisms mediating serial dependence effects across audition and vision, we first focused on the properties of this effect across the two modalities, at the behavioural level. Namely, we assessed two important characteristics of the effect: its selectivity for the features of the stimuli (“feature-selectivity” condition), and for their spatial position (“spatial-selectivity” condition).

Regarding the feature selectivity of serial dependence in different modalities, our prediction was that while it should be weak or absent in vision – in line with previous studies in magnitude perception (Fornaciai & Park, 2019b; Togoli et al., 2021), audition might instead show a more pronounced selectivity, especially when it comes to a fundamental and salient dimension like the pitch of a pure tone. Regarding the spatial selectivity of the effect, we instead predicted an opposite pattern of effects, in line with the intrinsic properties of audition and vision. Indeed, while vision entails a spatially localised encoding of the stimuli throughout its processing hierarchy, with a fine spatial resolution, the auditory modality is characterised by a poor spatial resolution (e.g., Alais & Burr, 2004). Previous results in vision indeed show some degree of spatial localisation of the serial dependence effect (Collins, 2019; C. Fischer et al., 2020; J. Fischer & Whitney, 2014; Fornaciai & Park, 2018b), but the same would hold true for audition is unknown. In Exp. 1 a total of 20 participants was included in the analysis.

In the experiment, participants compared the duration of a constant reference (200 ms) with a probe duration varying from trial to trial (100-400 ms). To induce attractive serial dependence effects, a task-irrelevant inducer stimulus (either 100 or 400 ms) was presented before the reference. The perceived duration of the reference as a function of the inducer and its properties (i.e., features or position) was assessed by fitting a cumulative Gaussian (psychometric) function to the proportion of responses at different probe levels. From the psychometric fit we extracted the point of subjective equality (PSE; the median of the fit, corresponding to chance level responses). Additionally, we also extracted the just noticeable difference (JND; the difference in duration between the 50% and 75% proportion of responses), which we used to compute the Weber’s fraction as a measure of precision in the task (WF = JND/PSE). In terms of serial dependence effect, what we expected is a relative underestimation of the reference duration when it was preceded by the shorter inducer, and a relative overestimation when it was preceded by the longer inducer. Then, to better assess the strength of the effect across conditions and modalities, we computed a serial dependence effect index based on the normalised difference between the PSE obtained with a 400-ms and a 100-ms inducer. In this context, a positive index shows the presence of an attractive effect between the inducer and the reference, while a negative index would show an opposite, repulsive effect.

The results of Exp. 1 are shown in Fig. 2A-B. Overall, what we observed is a clear double dissociation between the properties of the effect in audition and vision. Namely, in audition the effect was strongly selective for the features of the stimuli (Fig. 2A) – dropping to almost zero when inducer and reference had different pitch – but did not show any spatial selectivity (i.e., identical effect irrespective of the spatial position of inducer and reference). Conversely, in vision (Fig. 2B) the effect did not show any selectivity for the features (i.e., colour) of the stimuli, while it showed instead a selectivity for their spatial position. To better assess the significance of these difference, we performed a linear mixed effect model regression adding “modality” (vision vs. audition), “condition” (feature vs. spatial selectivity), and “congruency” (same vs. different, referring to either features or position) as fixed effects, and the subjects as random effect (R^2^ = 0.30). The results showed a significant three-way interaction between the different factors (β = -16.47, t = -2.37, p = 0.019), which was followed up with a series of one-sample and paired t-tests. First, with the exception of the auditory condition with different features (one-sample t-test against zero, t(19) = 1.21, p = 0.24; effect = 2.12%), all the effects in the individual conditions were significantly higher than zero (t(19) = 2.99-5.94, max p-value = 0.008; effects spanning from 7.62% in visual/different positions condition to 16.62% in the visual/same features condition). More importantly, to assess the selectivity of the effect, we compared the effect obtained with stimuli presented with the same features or same position vs. stimuli with different features or presented in different positions, in the two modalities. In audition, we observed a significant difference in the effect between stimuli with same and different features (t(19) = 3.15, p = 0.005, Cohen’s d = 0.84), while no difference was observed as a function of spatial position (same vs. different position; t(19) = 0.21, p = 0.84, d = 0.05). The opposite pattern was instead observed in vision. While the effect did not show any significant difference according to the congruency of the features (same vs. different features; t(19) = 0.54, p = 0.60, d = 0.11), it did show a significant difference according to the position of the stimuli (t(19) = 2.14, p = 0.045, d = 0.53). Taken together, these results show that serial dependence works differently in audition and vision, following a pattern of selectivity specific for the sensory modality tested.

**FIGURE 1.**
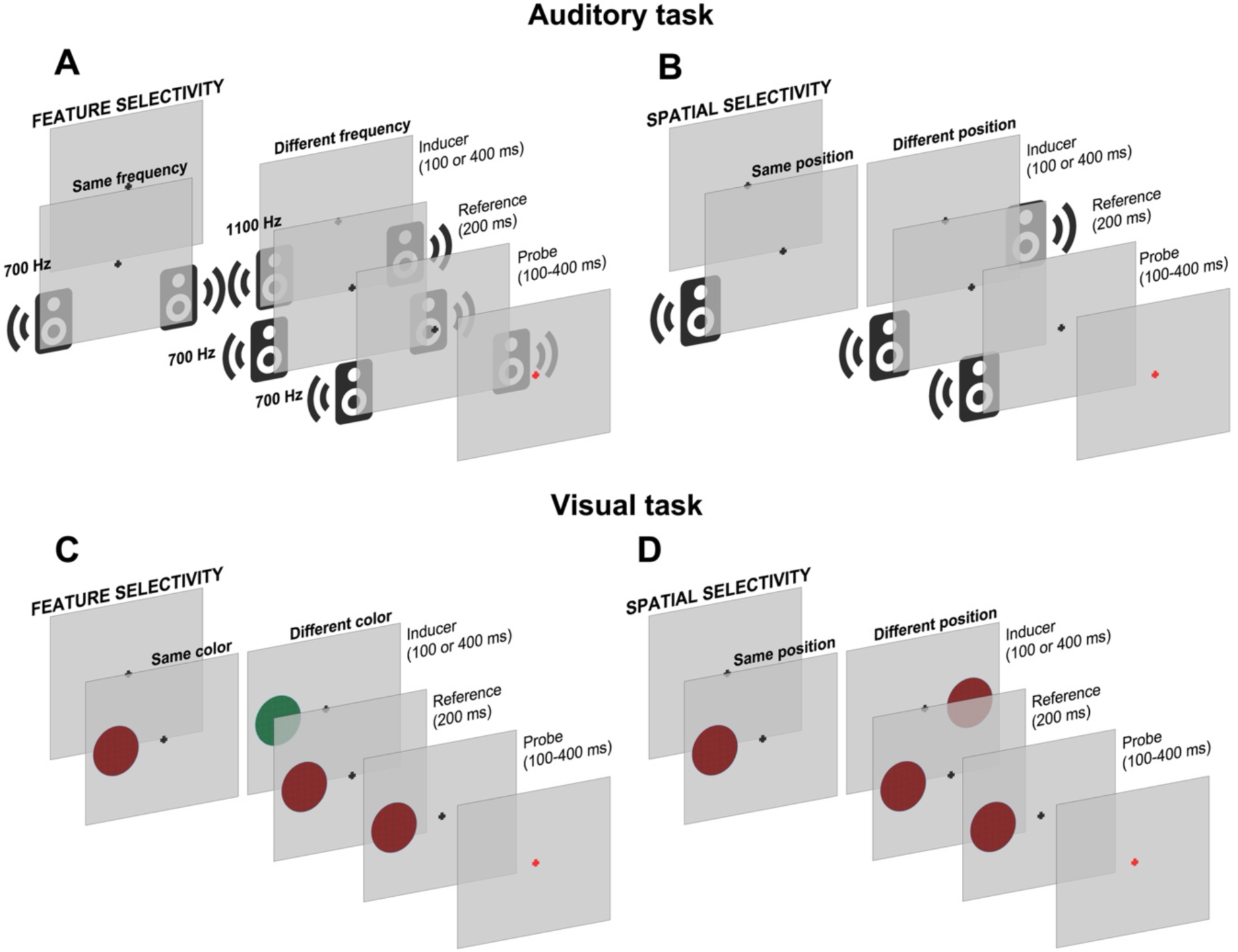
General paradigm employed in Exp. 1 and Exp. 2. (A-B) Procedure of the auditory duration discrimination tasks assessing the feature selectivity (A; “feature-selectivity” condition) and spatial selectivity (B; “spatial-selectivity” condition) of the attractive serial dependence effect. Three stimuli (pure tones) were presented in each trial: a task-irrelevant inducer (either 100 or 400-ms long) aimed to induce an attractive effect, followed by a constant reference (200 ms) and a variable probe (100-400 ms). In the feature selectivity condition, the inducer could have either the same pitch of the reference (700 Hz), or a different pitch (1100 Hz). In the spatial selectivity condition, the inducer could be presented either from the same speaker as the reference, or from a different speaker far enough to be clearly perceived as coming from a different source. At the end of each trial, participants reported which stimulus between the reference and probe lasted longer in time. (C-D) Visual duration discrimination tasks assessing feature selectivity (C) and spatial selectivity (D). The procedure was identical to the auditory task, with the exception that the stimuli were visual textures rather than auditory tones. In the feature-selectivity condition, the inducer could either be similar to the reference (same red and black texture), or different (green and black texture). In the spatial-selectivity condition, the inducer could be presented either in the same position as the reference, or in the opposite hemifield. Except for a few differences in the timing of the stimulus presentation, the paradigms of Exp. 1 and Exp. 2 were identical. In Exp. 1, the inter-stimulus interval (ISI) between the inducer and reference was 200-250 ms, while the reference-probe ISI was 350-450 ms. In Exp. 2 the ISI between the inducer and reference was 500-600 ms, while the reference-probe ISI was 800-900 ms. In both cases the inter-trial interval was 300-400 ms. Stimuli are not depicted in scale.

**FIGURE 2.**
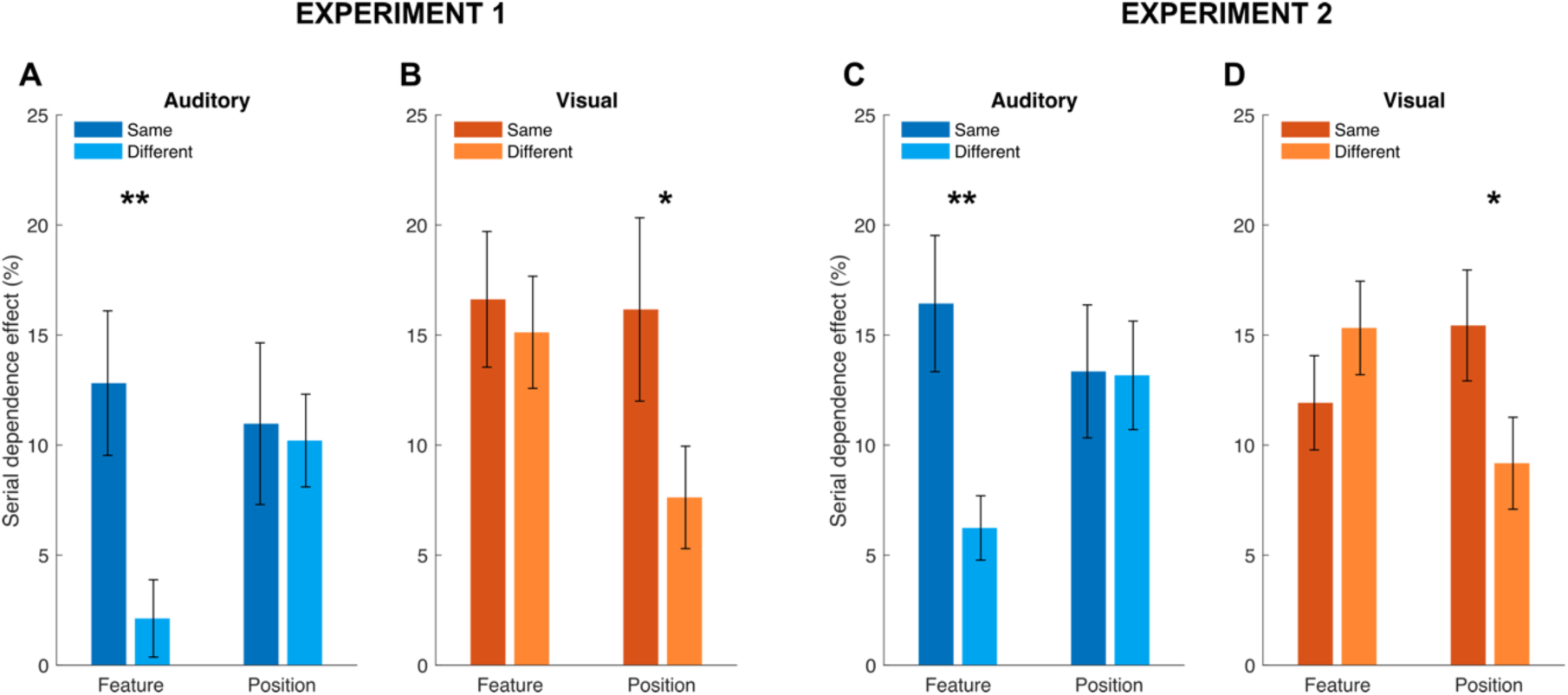
**Behavioural serial dependence effects measured in Exp. 1 and Exp. 2**. (A-B) serial dependence effects measured in audition (A) and vision (B) across the different conditions (feature-selectivity, spatial-selectivity) of Exp. 1, in terms of serial dependence effect index. (C-D) Serial dependence effects measured in Exp. 2. Stars refer to the significance of paired t-tests: * p < 0.05, ** p < 0.01. Error bars are SEM.

### Experiment 2

In Exp. 2, we used an identical paradigm compared to Exp. 1, but with the addition of EEG aiming at capturing and comparing the neural signature of perceptual history in audition and vision. Differently from the fully within-subject design of Exp. 1, here two independent groups of participants performed the feature-selectivity and spatial-selectivity condition. A total 23 participants were included in each condition.

First, the behavioural results obtained in Exp. 2 replicated the findings of Exp. 1. To assess the pattern of results across the two conditions (shown in Fig. 2C-D), we used again a linear mixed effect model analysis including, condition (as between-subject factor), congruency, and modality as fixed effects, and the subjects (coded independently for the two conditions) as random effect (R^2^ = 0.37). The analysis showed again a significant three-way interaction between condition, modality, and the congruency of features/positions (β = -19.68, t = -3.64, p < 0.001), which was followed up with individual one-sample t-tests and paired t-tests. The results from the series of one-sample t-tests showed significant serial dependence effects across all the conditions (t(22) = 4.27-7.21, all p-values < 0.001). The results of paired t-tests comparing the effect across in the same features/position vs. different features/position showed an identical pattern compared to Exp. 1. Namely, in audition (Fig. 2C), serial dependence resulted to be significantly stronger when inducer and reference shared the same pitch compared to when they had a different pitch (t(22) = 3.65, p = 0.001, d = 0.79), while no significant difference was observed when the stimuli were presented in the same or in a different position (t(22) = 0.07, p = 0.94, d = 0.01). The opposite was true in vision (Fig. 2D), where serial dependence did not significantly differ based on the features of the stimuli (same vs. different; t(22) = -1.35, p = 0.19, d = 0.33), but it was instead significantly stronger when inducer and reference were presented in the same position (t(22) = 2.39, p = 0.025, d = 0.56). Overall, these results, obtained in an independent group of participants, closely replicate the results obtained in Exp. 1, suggesting that this is indeed a robust pattern likely based on stable properties of serial dependence in these different sensory modalities.

Additionally, since we did not want to confuse the lack of spatial selectivity of the effect in audition with a trivial inability of participants to discern the different locations of the auditory stimuli (i.e., considering that auditory spatial localisation is notoriously poorer compared to vision; Alais & Burr, 2004), we also tested the ability of the participants to discriminate the location of the sound, by presenting it either from one speaker or the other in a brief session preceding the main experiment. Doing so, we confirmed that the different positions of the stimuli were clearly and consistently discriminable (accuracy ∼99%), suggesting that the lack of spatial selectivity of the auditory serial dependence effect is not due to the inability of participants to discriminate the different locations where the sound was presented.

### EEG results – feature-selectivity condition

Regarding the EEG results (Fig. 3) we first identified a set of target channels to perform further analyses. To select the target channels, we computed the difference in the responses evoked by the reference stimulus as a function of the duration of the preceding inducer, irrespective of its features, and plotted the topographic distribution of this signature of perceptual history across the scalp (see Fig. 3A). We then chose the channel showing the peak amplitude and a nearby channel showing the highest amplitude among the surrounding electrodes. We chose to use the average of two channels in the analysis (instead of just one) to improve the signal-to-noise ratio. Based on this procedure, in audition we selected the channels P8 an T8 in the right hemisphere. Activity averaged across these two channels showed a peak at 390 ms after stimulus onset (1.21 μV). The topography of average activity in a 50-ms window around this peak is shown in Fig. 3A. In vision, the contrast amplitudes showed a more posterior topography, peaking at channels P8 and PO4. The peak at these two channels was observed at a similar latency compared to audition (414 ms, 0.87 μV; Fig. 3A). Fig. 3B-C shows the average ERPs across the selected channels time-locked to the onset of the reference, separately for the two inducers duration. As shown in the figure, the duration of the preceding inducer had a noticeable impact on the amplitude of ERPs in both modalities, showing a marked upward shift in amplitude with increasing inducer duration.

**FIGURE 3.**
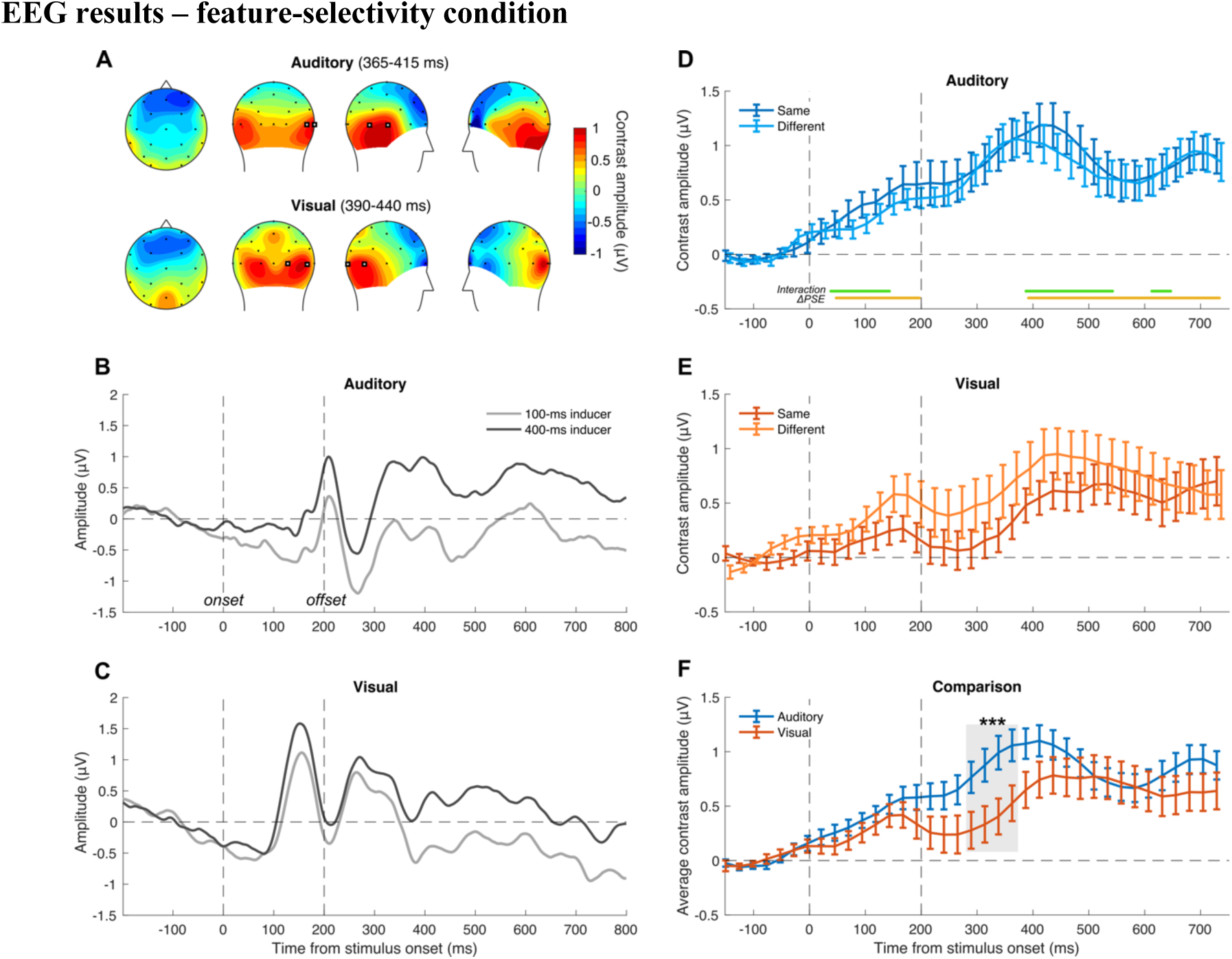
EEG results of the feature-selectivity condition of Exp. 2. EEG data was assessed by analysing activity in an epoch spanning from -200 to 800 ms around the onset of the reference stimulus, with the pre-stimulus interval used to perform baseline correction. EEG responses to the reference were binned separately according to the duration of the preceding inducer (100 ms or 400 ms) and according to its features (same or different), and averaged across trials to compute the event-related potentials (ERPs). (A) Topographic plots of activity across the scalp, in a 50-ms time window around the peak observed within each modality, used to select the target channels for further analyses. To do so, we computed the average difference in ERPs between the 100- and 400-ms inducer, irrespective of the features of the stimuli, which was taken as an overall measure of the effect of perceptual history on brain evoked responses. As target channels we selected the channel showing a peak in the contrast between different inducer conditions and the immediately adjacent channel showing highest amplitude among the surrounding electrodes. The chosen target channels, highlighted in the figure with bold markers, are T8 and P8 in audition, and P8 and PO4 in vision. (B) Auditory ERPs time-locked to the onset of the reference, plotted separately for the cases where the reference was preceded by a 100-ms or a 400-ms inducer, irrespective of the features of the stimuli. (C) Visual ERPs time-locked to the reference presentation, as a function of the inducer duration (again, irrespective of the features of the stimuli). (D) Contrast waves (i.e., difference between the ERPs evoked by the reference as a function of the inducer duration) in the auditory modality, in the same and different features conditions. The yellow and green lines at the bottom of the plot mark the significant clusters of time windows observed in the LME analysis. (E) Contrast waves in the visual modality. (F) Comparison of the contrast waves obtained in the auditory and visual modality, averaged across the same and different features condition. Note that the contrast waves shown in panels D-F have been smoothed with a sliding window average (100 ms window with 25 ms step). The shaded area highlights the time window showing a significant difference between audition and vision. The stars refer to the significance of a cluster-based non-parametric test, *** p < 0.001. Error bars are SEM.

To better assess the neural signature of perceptual history in the two modalities, we focused on the difference (“contrast”) in ERP amplitude as a function of the inducer duration, measured according to whether inducer and reference had the same or different features. To further increase the signal-to-noise ratio we computed the average contrast amplitude across a series of 100-ms windows (with 25 ms step). Additionally, to control for multiple comparisons, we opted for a non-parametric approach, first setting a cluster-size threshold of at least 3 consecutive significant time points and then performing a cluster-based permutation test. The contrast amplitude in audition (Fig. 3D) showed an increasing pattern starting from stimulus onset and peaking at around 370-410 ms (1.18 μV and 1.05 μV, respectively for the same and different features condition). The pattern was almost identical across the same and different features condition, and did not show any significant difference (paired t-tests across conditions, max t(22) = 1.12, min p = 0.10). In vision (Fig. 3E), the contrast amplitude showed a different pattern, with an earlier peak at 150-170 ms (0.25 μV and 0.60 μV for the same and different features condition), followed by a later, higher peak at around 450-510 ms (0.65 μV and 0.94 μV). In this case, we observe a few isolated windows showing a significant difference across the same and different condition in the pre-stimulus interval (-150 and -50:-25 ms; max t(22) = -2.17, min p = 0.041), but none of these exceeded the cluster-size threshold. Interestingly, the contrast amplitude in vision seemed mostly higher (although not significantly) when inducer and reference had different features compared to when they were similar, potentially in line with the slightly stronger behavioural serial dependence effect observed in this condition. However, in the auditory modality – which is where we observed the strongest difference in serial dependence as a function of the features of the stimuli – we did not observe any difference in the neural effect of the inducer as a function of its features.

In order to characterise the link between the behavioural and neural signature of perceptual history, we used a linear mixed-effect (LME) regression model performed across each time window within the reference epoch. Namely, we assessed the relationship between the difference in PSE as a function of the inducer (ΔPSE = PSE_400ms_ – PSE_100ms_) and the amplitude of the contrast wave (ΔERP), adding the features (same vs. different) of the stimuli as an additional fixed effect factor and the subjects as the random effect (βPSE ∼ βERP × Features + (1|subj)). Again, to control for multiple comparison, we set a cluster-size threshold of at least three consecutive significant windows, and tested the significance of each individual cluster with a non-parametric cluster-based test (see *Methods*). The clusters showing a significant relationship between the behavioural (ΔPSE) and neural (ΔERP) signature of perceptual history are marked with yellow lines in Fig. 3D-E, while the significant interaction between ΔERP and the features of the stimuli is marked with green lines. In audition, we observed two main clusters of significant time windows for the effect of ΔERP: one early cluster spanning from 50 to 200 ms (seven consecutive windows, t ≥ -2.29, p ≤ 0.02, cluster p < 0.001), and a later one spanning from 380 to 730 ms (15 consecutive windows, t ≥ -2.27, p ≤ 0.03, cluster p < 0.001). More interestingly, we observed three clusters of time window where the serial dependence effect is significantly predicted by the interaction between ΔERP and the features of the stimuli: 40-140 ms (five consecutive windows, t β 2.21, p β 0.03, cluster p < 0.001), 380-540 ms (six consecutive windows, t β 2.13, p β 0.04, cluster p< 0.001), and 610-650 ms (three consecutive windows, t β 2.07, p β 0.044, cluster p < 0.001). Conversely, however, in vision we did not observe any significant relationship between the behavioural serial dependence effect and the difference in ERP amplitude caused by the inducer, nor a significant interaction between the ΔERP and the features of the stimuli.

These results show that at least in audition the strength of the serial dependence effect is significantly related to the impact that the inducer has on visual evoked potentials, and importantly, that such relationship emerges quite early in the visual processing stream, starting at around 50 ms after stimulus onset. In vision, however, this relation could not be captured by the linear regression model.

Finally, we compared the average contrast amplitude across the two modalities, to assess whether this signature shows any significant difference across the time-course of the reference processing (Fig. 3F). In this context, we observed that the contrast amplitude is overall lower in the visual modality, and a significant difference between vision and audition could be observed in a cluster spanning from 280 to 360 ms (paired t-tests, four consecutive significant windows, t β 1.83, p β 0.01, cluster p < 0.001).

Furthermore, in order to potentially capture different effects that cannot be captured with a linear model, we performed a non-linear regression analysis. To perform this analysis, we considered the trial-by-trial EEG activity rather than the averaged ERPs. Namely, we used the EEG activity in each trial (averaged across a series of 100-ms windows with a 25-ms step) as dependent variable. As predictors, we entered the response in each trial (0 or 1, corresponding to “reference longer” or “probe longer”), the duration of the inducer (100 ms or 400 ms), and the features of the stimuli (same or different). The resulting beta values for each of these factors thus provide a measure of whether and to what extent they modulate the EEG responses across individual trials. Note that while the inducer duration and the features of the stimuli reflect properties that are already evident at the onset of the reference, the response will only occur much later in the trial, after the presentation of the probe stimulus. This allows to exclude the possible confound of motor preparation, since seeing the reference itself is not enough for participants to start preparing the response and the related motor action. The results of this analysis in the auditory modality are shown in Fig. 4A. To assess the significance of beta values, we performed a series of one-sample t-tests against zero, and again performed a cluster-based non-parametric test after applying a cluster-size threshold of at least three consecutive significant time windows. First, regarding the influence of the inducer on EEG responses, we observed two main clusters of significant time windows: 70-140 ms (four consecutive time windows, t ≥ 2.09, p β 0.02, cluster p = 0.0006), and 220-530 ms (14 time windows, t ≥ 2.14, p β 0.044, cluster p < 0.001). Regarding the influence of the features of the stimuli, we observed two significant clusters at 240-290 ms (three windows, t ≥ -2.19, p ≤ 0.04, cluster p < 0.001) and 410-680 ms (12 windows, t ≥ -2.19, p ≤ 0.04, cluster p < 0.001). Finally, regarding the relationship between EEG activity and the behavioural response, we observed again two clusters spanning 310-490 ms (eight windows, t ≥ 2.51, p β 0.02, cluster p < 0.001), and 580-680 ms (five windows, t ≥ -2.10, p ≤ 0.047, cluster p < 0.001). This shows again that perceptual history and the features of the stimuli strongly affect EEG responses even when considered in a trial-by-trial fashion, and that brain activity shortly after the reference presentation is related to the choice that the participants will make much later in the time-course of a trial. Regarding the visual condition, we did not observe any consistent effect of the inducer duration and the congruency in the features of the stimuli on the trial-by-trial EEG activity. However, we found again a significant effect of the response, at a single cluster of time windows spanning from 290 to 430 ms (seven windows, t ≥ 2.45, p β 0.02, cluster p < 0.0001) – which is very similar to the latency of this effect in the auditory modality. This shows that the association between behavioural responses and brain activity during the presentation of the reference can be observed irrespective of sensory modality at a similar time window.

**FIGURE 4.**
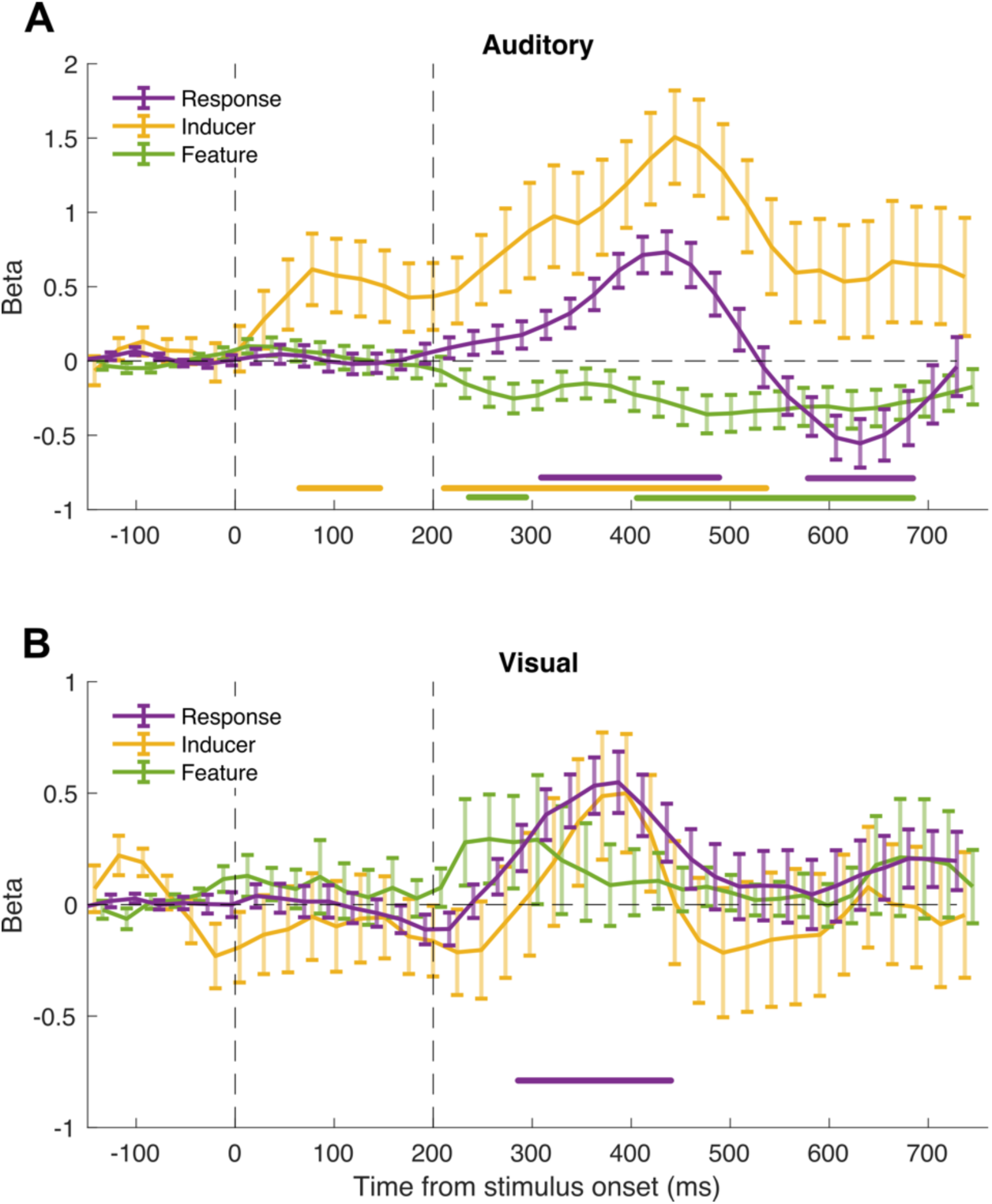
Results of the non-linear regression analysis in the feature selectivity condition. The non-linear regression analysis was performed by considering the trial-by-trial EEG responses, averaged across a series of 100-ms time windows across the reference epoch, as a function of the duration of the inducer, the similarity/difference of inducer and reference features, and the response that the participants will provide at the end of the trial. The beta values obtained in this analysis represent the extent to which the EEG amplitude is modulated by the different levels of the different factors. For example, a beta value of 0.5 means that the amplitude of EEG responses increases by 0.5 μV across the levels of the factor. (A) Results of the non-linear regression analysis in the auditory modality, in terms of average beta values across the group, at each time window throughout the analysed epoch. (B) Results of the regression analysis in the visual modality. The lines at the bottom of each panel indicate the significant clusters observed in the analysis, with the colour matching that of the main plots. Error bars are SEM.

### EEG results – spatial-selectivity condition

After assessing the properties of the neural signature of perceptual history in the feature-selectivity condition and its relationship with the behavioural serial dependence effect, we went on and performed a similar set of analyses on the data from the spatial-selectivity condition.

First, in the spatial-selectivity condition, ERPs were computed in a slightly different way compared to the feature-selectivity condition, to account for the different positions of the stimuli. Namely, the final ERPs were computed as the average of the channels contralateral to the reference stimulus positions – i.e., the average of left channels (T7, P7) in the trials where the reference was presented on the right of the fixation point, and the right channels (T8, P8) when the reference was presented on the left. The topographic distribution of activity seemed indeed to be localised to electrodes contralateral to the reference stimulus, especially in the visual condition (Fig. 5A). In audition this lateralisation was less evident, but we implemented the same strategy for consistency with the visual condition. The overall peak of activity in the visual modality was observed at 95 ms after stimulus onset (0.45 μV), while at about 400 ms (0.71 μV) in the auditory modality. Fig. 5B-C shows examples of the ERPs time-locked to the reference stimulus, sorted according to the duration of the inducer stimulus (100 and 400 ms), but irrespective of the position of the stimuli.

**FIGURE 5.**
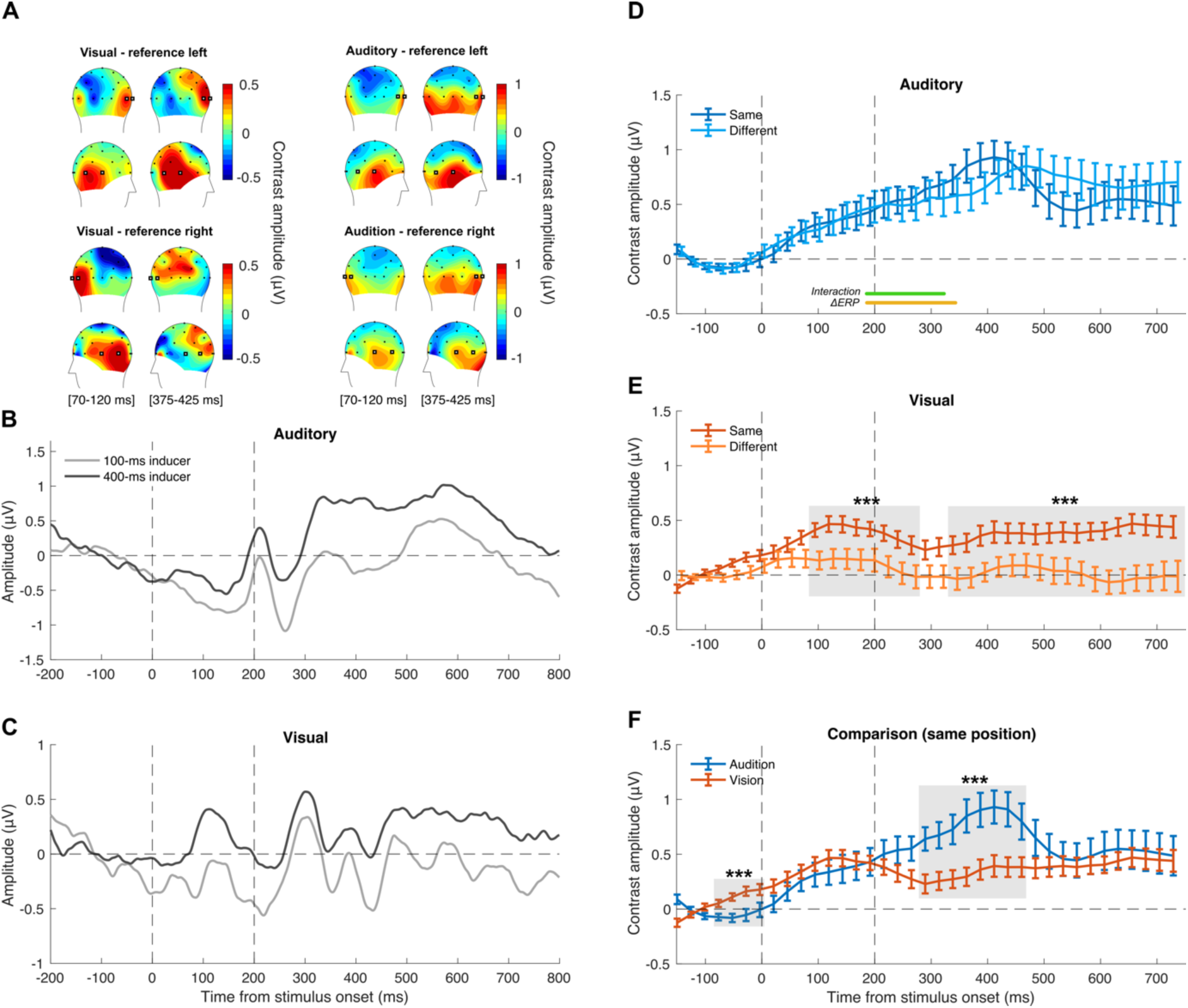
EEG results of the spatial-selectivity condition of Exp. 2. (A) Topographic plots of activity across the scalp, across two 50-ms time windows around the peaks observed in the visual (70- 120 ms) and auditory (375-425 ms) modality, which were used to select the target channels for further analyses. In this case, we selected a set of two electrodes similar across the two modalities: T8 and P8 in the right hemisphere, and T7 and P7 in the left hemisphere. The selected channels are highlighted in the figure with bold markers. To capture the spatially-selective effect according to the different positions of the stimuli, ERPs were computed considering the average of electrodes contralateral to the position of the reference stimulus. (B) Auditory ERPs time-locked to the onset of the reference, plotted separately for the cases where the reference was preceded by a 100-ms or a 400-ms inducer, irrespective of the position of the stimuli. (C) Visual ERPs time-locked to the reference presentation, as a function of the inducer duration. (D) Contrast waves (i.e., difference between the ERPs evoked by reference as a function of the inducer duration) in the auditory modality, in the same and different position conditions. The yellow and green lines at the bottom of the plot mark the significant clusters of time windows observed in the LME analysis. (E) Contrast waves in the visual modality. (F) Comparison of the contrast waves obtained in the auditory and visual modality, averaged across the same and different position condition. Contrast waves shown in panels D-F have been smoothed with a sliding window average (100 ms window with 25 ms step). The shaded area highlights the time window showing a significant difference between audition and vision. The stars refer to the significance of a cluster-based non-parametric test. Error bars are SEM. *** p < 0.001.

Similarly to the feature-selectivity condition, here we focused on the contrast between different inducer durations according to the position of inducer and reference stimuli (same vs. different; Fig. 5D-F). In audition (Fig. 5D), we observed a pattern largely similar to the feature-selectivity condition, with the contrast amplitude – the signature of the inducer effect – increasing from stimulus onset up until around 400 ms. Comparing the same vs. different condition with paired t-tests across all the time windows showed no significant difference across them (max t(22) = 1.21, min p = 0.22). In vision, instead, the congruence of inducer and reference position had a strong impact on the effect of the inducer at the neural level. Indeed, we observed a much stronger difference in the response evoked by the reference as a function of the preceding inducer stimulus when the two occupied the same spatial position. Running a series of paired t-tests (complemented with a cluster-based non-parametric test and a cluster-size threshold of at least three significant windows) showed two main clusters of significant time windows spanning almost the entire epoch: 90-270 ms (eight consecutive windows, t β 2.11, p β 0.04, cluster p < 0.001), and 340-750 ms (17 windows, t β 2.17, p β 0.048, cluster p < 0.001). Moreover, to address the link between the behavioural and neural signature of perceptual history, we performed again a linear mixed-effect regression analysis at each time window throughout the epoch (ΔPSE ∼ ΔERP × Position + (1|subj)). Again, we used the difference in PSE observed as a function of the inducer duration (ΔPSE) as dependent variable, and assessed the extent to which the behavioural effects across the group could be predicted by the difference in amplitude as a function of the inducer duration (i.e., the amplitude of the contrast wave, ΔERP) and its interaction with the position of the stimuli (same vs. different). In audition (Fig. 5D), we observed one cluster of significant effect of ΔERP, spanning from 190 to 340 ms (seven windows, t ≥ -2.37, p ≤ 0.02, cluster p < 0.001). Overlapping with this cluster, we also observed a significant interaction between ΔERP and the positions of the stimuli (190-320 ms, six windows, t ≥ 2.04, p β 0.047, cluster p < 0.001), even if the position of the auditory stimuli did not seem to affect either the behavioural effect or the ERPs. On the other hand, in vision (Fig. 5E) we did not observe any significant relationship between the behavioural and neural effect, similarly to the results of the feature-selectivity condition. Finally, we also compared the contrast waves of the two sensory modalities against each other (limited to the case where inducer and reference were presented in the same position), to further assess the difference in the dynamics of this signature across vision and audition (Fig. 5F). Similarly to the feature-selectivity condition, we found that the amplitude of the contrast wave differs across modalities at around 290-460 ms (eight consecutive windows, t ≥ -2.18, p ≤ 0.03, cluster p < 0.001). In addition to this cluster, we also observed an earlier and smaller cluster before stimulus onset, from -80 to 0 ms (four windows, t ≥ 2.17, p β 0.03, cluster p < 0.001). This may suggest that some signature of perceptual history may be evident even before the onset of the reference stimulus, at least in vision.

Again, in vision the relationship between the neural signature of perceptual history and the behavioural serial dependence effect has proven to be more difficult to capture compared to the auditory modality, at least with a linear regression analysis performed on measures such as the PSE and the ERPs. We thus performed again a non-linear regression analysis. In this analysis we entered as dependent variable the EEG responses, averaged across different time windows, measured in each individual trials. As predictors, we used the inducer duration (100 vs. 400 ms), the position of the stimuli on the screen (left vs. right hemifield), the spatial match of inducer and reference position (same vs. different), and the response that the participant will provide at the end of the trial (0 or 1, i.e., “reference longer” or “probe longer”).

The results of this non-linear regression analysis are shown in Fig. 6. In the auditory modality (Fig. 6A), consistently with the results of the feature-selectivity condition, we found a significant relationship between EEG amplitude and the participants’ response in the task, at a window spanning 360-490 ms (six windows, t ≥ 2.41, p β 0.02, cluster p < 0.001). The later negative deflection in beta values however did not reach significance in this condition. Besides that, we observed an effect of the inducer duration at around 410-460 ms (three windows, t ≥ 2.08, p β 0.049, cluster p < 0.001). No significant influence of the spatial match between inducer and reference, but, surprisingly, we observed a small but significant effect of position (left vs. right). This effect indicates that EEG amplitude differed slightly, but systematically, according to the position of the reference irrespective of whether it was consistent with the inducer or not. The effect of position was evident across three clusters, spanning -5-70 ms (t β 2.31, p β 0.03, cluster p < 0.001), 240-410 ms (t β 2.09, p β 0.048, cluster p < 0.001), and 570-730 ms (t β 2.30, p β 0.03, cluster p < 0.001). Finally, in the visual modality, we observed again a significant relationship between EEG and behavioural responses, in a cluster of time windows spanning from 290 to 460 ms (eight windows, t β 2.20, p β 0.04, cluster p < 0.001). In line with the spatial selectivity of the effect, we also observed two significant clusters showing an effect of spatial match on EEG activity, at 240-290 ms (three windows, t ≥ -2.15, p ≤ 0.04, cluster p < 0.001), and 480-530 ms (three windows, t β 2.18, p β 0.04, cluster p < 0.001).

**FIGURE 6.**
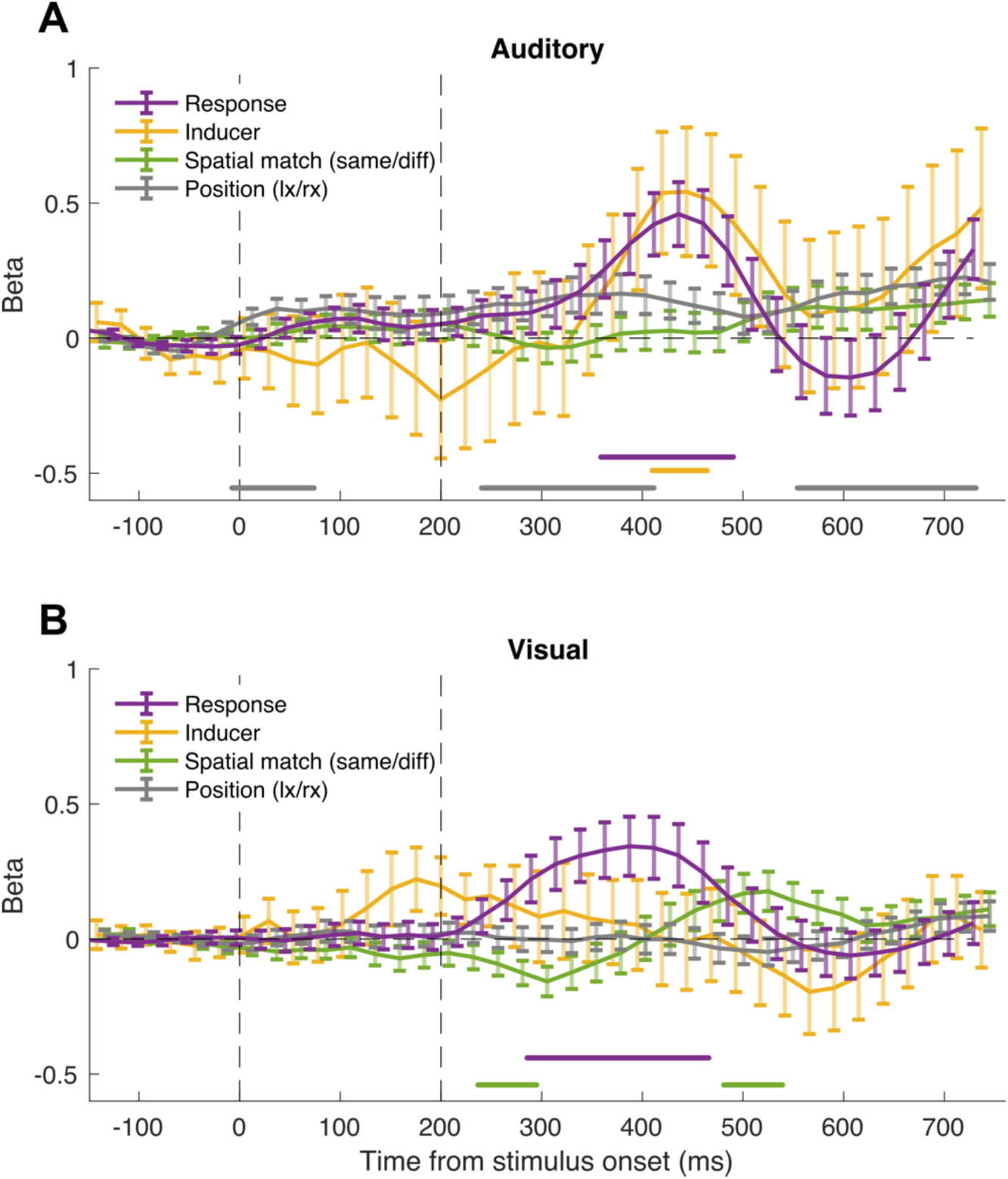
Results of the non-linear regression analysis in the spatial-selectivity condition. (A) Results of the non-linear regression analysis in the auditory modality, in terms of average beta values across the group, at each time window throughout the analysed epoch. (B) Results of the regression analysis in the visual modality. The lines at the bottom of each panel indicate the significant clusters observed in the analysis, with the colour matching that of the main plots. Error bars are SEM.

## DISCUSSION

In this study, we aimed to disentangle the mechanisms mediating the influence of perceptual history across different sensory modalities. Perceptual history has been shown to play an important role in perception, shaping how we experience the external stimuli according to the recent history of stimulation. Indeed, attractive “serial dependence” effects, especially in visual perception, have started to attract increasing interest in the past decade due to the widespread and systematic way in which these effects emerge in perceptual tasks. In serial dependence, a stimulus that we are currently seeing is attracted by the stimuli in the recent past, effectively appearing more similar to the preceding stimuli than it actually is in reality. A growing number of studies show that serial dependence is ubiquitous in vision (Alais et al., 2017; J. Fischer & Whitney, 2014; Fornaciai & Park, 2018b; Liberman et al., 2014), potentially suggesting that exploiting perceptual history is a general computational principle characterising perception and decision-making as a whole. In audition, evidence for the existence of serial dependence effects is more limited, but attractive influences of perceptual history have been reported also in this modality (e.g., Motala et al., 2020).

Considering this seemingly generalised influence of perceptual history across different dimensions and modalities, our question thus is: is the effect of perceptual history mediated by a common, perhaps centralised, processing mechanism, shared by different perceptual domains and sensory modalities? Or is it the result of similar computations implemented independently across different perceptual pathways?

So far, the neural mechanisms mediating the attractive effect of perceptual history are poorly understood. According to a recent framework proposed to explain serial dependence (e.g., Burr & Cicchini, 2014; J. Fischer & Whitney, 2014; Manassi & Whitney, 2022), this attractive bias would reflect the operation of a perceptual stability mechanism smoothing and filtering out noise from sensory signals. According to this idea, perceptual history would thus operate at the level of perceptual processing, effectively modulating the phenomenological appearance of a stimulus. Alternatively, serial dependence has been interpreted as a more “cognitive” effect, occurring at the level of perceptual decision-making (Pascucci et al., 2019; Wehrman et al., 2020) or during working memory encoding (Bliss et al., 2017; but see Manassi et al., 2018). Although behavioural results suggest that serial dependence is generated at a high-level of the brain processing hierarchy – as shown for instance by its dependence on conscious perception and feedback processing (Fornaciai & Park, 2019a, 2021) – neuroimaging results nevertheless support the idea that perceptual history operates at a relatively low perceptual level. For instance, fMRI results have shown that the primary visual cortex (V1) seems involved in serial dependence in orientation perception (st. John-Saaltink et al., 2016). Other studies employing the EEG technique further support the involvement of early sensory/perceptual processing stages in mediating the effect of serial dependence, showing a signature of perceptual history emerging early in brain signals (50-200 ms; Fornaciai & Park, 2018a, 2020), and hence supporting the idea that perceptual history shapes the appearance of a stimulus, not just how we judge or remember it.

If perceptual history truly operates at the level of perceptual processing, then we predict the existence of at least partially independent mechanisms operating before distinct sensory pathways converge onto associative brain areas. To address this possibility, we focused on time perception, and assessed the behavioural and neural signature of perceptual history effects in a duration discrimination task (Togoli et al., 2021). Time perception was specifically chosen since, differently from other perceptual domains more often studies in this context (i.e., orientation or numerosity perception), interval duration can similarly be delivered as a visual or an auditory input, which is crucial in order to compare the results across different modalities. Our results, overall, provide evidence that this is indeed the case, by demonstrating that (1) serial dependence works differently in different sensory modalities, and that (2) perceptual history shows at least partially different neural signatures in vision and audition.

First, the behavioural results of Exp. 1 and Exp. 2 show a clear double dissociation in the properties of serial dependence in audition and vision. Namely, while serial dependence in audition shows a strong selectivity for the features (i.e., pitch) of the stimuli – yielding a significantly stronger effect when successive stimuli are similar to each other – it is not very selective for the position of the stimuli, even if they are presented in clearly different and discriminable positions. Conversely, vision does not show any selectivity for the features (i.e., colour) of the stimuli, while it instead shows a significantly stronger effect when the stimuli are presented in the same position. Interestingly, these results are consistent with the “specialisation” of different sensory modalities to different perceptual dimensions. Indeed, while human vision has a fine spatial resolution but poor sensitivity to time, the opposite is true for audition, which shows a much finer temporal resolution and poor spatial resolution (e.g., see for instance Alais & Burr, 2004; Chen & Yeh, 2009; Heron et al., 2013). Regarding the effects in vision, the fact that some effect could be observed even when the stimuli are presented in different positions is consistent with the relatively broad spatial localisation of the effect shown in previous studies (Collins, 2019; J. Fischer & Whitney, 2014; Fornaciai & Park, 2018b). This in turn suggests that the effect occurs within a topographically organised map of the visual environment, but that it also involves either very large receptive fields, or feedback signals from mid- or high-level areas. The possibility of the involvement of feedback signals is consistent with previous observations of a crucial role of feedback processing in serial dependence (Fornaciai & Park, 2019a, 2021). Moreover, the tolerance of the effect for differences in the features of the stimuli is consistent with previous results in numerosity perception (Fornaciai & Park, 2019b, 2022), although in other domains (i.e., orientation perception) the effect has been shown to be more selective for the contextual features of the stimuli (C. Fischer et al., 2020). Overall, this different pattern of effects suggests that modality-specific processing mechanisms, working according to the intrinsic properties of different modalities, likely mediate the behavioural effect of perceptual history. Finally, the existence of independent mechanisms involved with perceptual history is also consistent with results showing that serial dependence does not work cross-modally (i.e., the numerosity of an auditory stimulus does not affect the perceived numerosity of a visual stimulus; Fornaciai & Park, 2019b), and, even within the same modality, does not work across different but related dimensions such as time and numerosity (Togoli et al., 2021).

Regarding the interpretation of EEG results – highlighting the neural signature of perceptual history – our findings show different patterns of activity that support the idea of different modality-specific mechanisms. As a first signature of perceptual history at the neural level, we considered the contrast between the ERPs evoked by the reference when it was preceded by the long (400 ms) versus the short (100 ms) inducer. Indeed, the dynamics and layout of individual ERPs (i.e., see Fig. 3B-C and Fig. 5B-C) is mostly determined by the different processing of auditory and visual stimuli, making it difficult to discern a difference purely based on the influence of perceptual history per se. Contrasting the difference in evoked responses as a function of the preceding inducer should instead provide a clearer measure of the impact of perceptual history on brain processing.

A first difference in the neural signature of perceptual history across modalities is the difference in the scalp topography of activity. Indeed, in the feature-selectivity condition, the contrast across different inducers had a more posterior distribution compared to audition, which instead showed more lateral peaks. In the spatial-selectivity condition, although we selected the same electrodes as target channels in both modalities, we observed a more bilateral distribution of activity in audition, while vision showed more localised foci contralateral to the reference stimulus. This difference, although interesting, is however limited by the poor spatial resolution of scalp EEG, especially considered that we used only 32 electrodes, and hence it must be interpreted with caution.

The dynamics of contrast waves at the selected target channels and the relation between the contrast amplitude and the behavioural effect provide instead clearer evidence for the nature of the perceptual history mechanisms implemented in different modalities. First, in both modalities the contrast amplitude starts to ramp up very early after stimulus onset, suggesting that a neural signature of perceptual history is present from the earliest stages of sensory processing – which is in line with previous EEG studies on serial dependence (Fornaciai et al., 2020; Fornaciai & Park, 2019a, 2020a). Second, however, the contrast amplitude unfolds differently in time, showing different peaks and a different overall strength. Specifically, while in audition the contrast amplitude increases up until a peak at around 400 ms, it shows a much earlier peak in the visual modality (∼150 ms). After its initial peak before stimulus offset, the contrast amplitude in vision tended indeed to decrease, leading to a significant difference compared to audition at around 300-400 ms. This pattern might in turn suggest a difference in the way perceptual history is propagated during stimulus processing, affecting different stages. For instance, the earlier peak observed in vision might suggest a faster integration of perceptual history into current sensory processing occurring during the stimulus presentation, while the increasing pattern observed in audition might suggest a slower and more sustained process.

Interestingly, while the spatial selectivity of the effect is well captured by the difference in contrast amplitude observed in the visual condition (Fig. 3E), the feature selectivity of the auditory effect does not seem to be reflected by the contrast amplitude, which unfolds similarly irrespective of the stimulus features. This shows that even in audition, a mismatch in the stimulus features does not necessarily prevent or reduce the encoding of perceptual history like a difference in position does in vision, suggesting an additional difference in how the selectivity is achieved in different modalities. Namely, while spatial selectivity could arise from a localised encoding of perceptual history, the feature selectivity observed in audition might stem from a mechanism preventing the influence of perceptual history when the past stimulus does not match the current one, even if perceptual history itself does remain encoded in brain responses (see also (Fornaciai et al., 2020) for a similar interpretation of the effects across different dimensions).

The results of the linear regression analysis show a significant link between the contrast amplitude measured with EEG and the behavioural effect, at least in audition. In the feature-selectivity condition, we indeed observed that the contrast amplitude in the auditory modality can successfully predict the behavioural serial dependence effect at early latencies, starting at around 50 ms after stimulus onset. In the spatial-selectivity condition, instead, this significant relationship seems to emerge right after the offset of the stimulus. This in turn suggests a difference in how the influence of perceptual history unfolds over time as a function of how the stimuli are manipulated. A possibility explaining this difference may be the predictability of the sequence of events in the two conditions. Namely, while in the feature-selectivity condition there was no uncertainty about the properties of the reference stimulus – which was always the same – the position of the reference in the spatial-selectivity condition was randomised. Thus, with more predictable stimuli, it is possible that perceptual history could start to play its role at a much earlier stage compared to less predictable stimuli.

On the other hand, in the visual modality, the linear regression model did not yield any significant effect, neither in the feature-selectivity condition, nor in the spatial-selectivity condition, despite the contrast amplitude showing a clear influence of the inducer on the brain responses evoked by the reference. We interpret this lack of a significant relationship between evoked activity and the behavioural effect – which was instead consistently captured in audition – as possibly another difference across the two modalities: while the behavioural and neural signature of perceptual history show a more direct relationship in audition, much easier to capture in data analysis, visual signals did not show such a clear relationship. If a shared mechanism was involved in generating the effect of perceptual history, then we would have expected to find comparable results across modalities. This, although speculatively, may thus highlight another specific property of the perceptual history mechanisms in vision and audition.

Although so far what we observed is more consistent with at least partially independent mechanisms implemented in different sensory modalities, the non-linear regression analysis showed instead a similarity between vision and audition. Specifically, we observed a significant relationship between the pattern of EEG activity across trials, and the behavioural (binary) response provided by participants, at similar latencies across modalities and experimental conditions. Indeed, this analysis shows that EEG activity in a broad window spanning from 300 to 500 ms after stimulus onset can be predicted, at least to some extent, by the response that the participant will provide at the end of the trial. Considering that the response will only be provided much later compared to the reference presentation, and the fact that the reference alone did not provide sufficient information for decision-making, what does this effect mean? Besides the intrinsic variability in stimulus representation that characterise perception, the only consistent source of variability that could affect participants’ choices is the effect of perceptual history. We thus interpret this result as a possible signature of a perceptual decision-making stage affected by perceptual history, which seems to be shared across modalities.

Interestingly, in all but one condition (i.e., visual spatial-selectivity), this effect peaked at around the same time of the inducer effect (i.e., as indexed by beta values reflecting the inducer duration in the non-linear analysis), which supports the idea that perceptual history may be responsible for this relationship between evoked responses and participants’ choices. In this context, we can speculate that the association with the behavioural response might represent a high-level processing stage shared by both vision and audition. For example, high-level cortices such as the supplementary motor area (SMA) might represent a possible candidate to support perceptual decision-making in time perception (Nani et al., 2019; Protopapa et al., 2019). Following the initial peak similar in vision and audition, the relationship with participants’ choice continues to unfold differently in the two modalities, showing a subsequent (opposite) peak at around 600 ms in audition, but no such later effect in vision.

To summarise, our results show three main differences in the signatures of perceptual history across audition and vision. First, the double dissociation in the properties of the behavioural effect provides evidence that serial dependence is governed by different “rules” consistent with the intrinsic properties of different sensory modalities. Second, the contrast of activity evoked by the reference as a function of the preceding inducer shows a different scalp topography (i.e., more posterior or more localised in vision, more temporal and bilateral in audition), and a different dynamic in the two modalities, with a more prominent peak before stimulus offset in vision, and a later, higher peak in audition. Third, in audition we observed a clear relationship between the behavioural and neural signature of perceptual history, which we were instead unable to capture in vision, suggesting a difference in how perceptual history is encoded in brain signals and how these signals relate to the behavioural effect. Using a non-linear regression model, on the other hand, we also highlighted a similarity between the two modalities: the relationship between EEG activity and choice in the task observed in a broad latency window similar across modality. Based on these data alone it is difficult to confidently conclude that such relationship stems from the exact same processing stage and neural generator, but its consistency across conditions at least suggest the possibility of a shared component, possibly related to perceptual decision-making.

A limitation that we need to acknowledge, however, is that our results are limited to the mechanisms of perceptual history in time perception, and hence the same properties demonstrated here may not generalise to different perceptual dimensions. Previous results on serial dependence indeed show different patterns when the effect is measured in different domains. For instance, in vision, while serial dependence in orientation perception is tuned to the similarity between successive stimuli and decreases when the stimuli become too different from each other (J. Fischer & Whitney, 2014; Pascucci et al., 2019), the effect in numerosity perception is more linearly related to the difference across stimuli (Fornaciai & Park, 2020b). The effect in orientation perception is also more sensitive to the contextual features of the stimuli (C. Fischer et al., 2020), while in numerosity (and also in time, as shown in the present work) visual serial dependence is more abstracted from the features of the stimuli (Fornaciai & Park, 2019b, 2022). However, these differences also suggest that serial dependence may involve partially independent mechanisms across different perceptual domains even within the same modality, making the existence of a completely generalised perceptual history mechanism less likely.

Overall, our results provide evidence that perceptual history in time perception operates differently in different sensory modalities, pointing to the existence of different, at least partially independent mechanisms implemented within different sensory pathways. The neural signature emerging early in ERPs also support the idea that irrespective of where perceptual history signals originate from in the brain (i.e., whether it is a low or a high level; Ceylan et al., 2021; Fornaciai & Park, 2019a, 2021), they operate at the level of perception. Finally, the existence of independent sensory modality-specific mechanisms would also be in line with a role of perceptual history in facilitating perceptual stability and continuity (J. Fischer & Whitney, 2014; Manassi & Whitney, 2022). To conclude, our results suggest that rather than being implemented in a centralised hub, the role that perceptual history plays in our conscious experience of the external world may stem from the same computational principle emerging independently in different brain processing pathways.

## METHODS

### Participants

A total of 64 participants took part in the study, with 20 subjects tested in Exp. 1 (14 females, age ± SD = 24.6 ± 2.4 years), and two groups of 25 subjects each tested in Exp. 2 (37 females, age ± SD = 25.1 ± 4.3 years). Two participants in the feature-selectivity condition of Exp. 2 were excluded from data analysis due to excessively poor performance (see *Behavioural data analysis*), while two participants were excluded from the spatial-selectivity condition of Exp. 2 due to equipment failure (i.e., EEG was not recorded properly). All participants reported normal or corrected-to-normal vision, and were naïve to the purpose of the study. Prior to taking part in the experiment, all participants read and signed a written informed consent form. All experimental procedures were approved by the ethics committee of the International School for Advanced Studies (SISSA) and were in line with the declaration of Helsinki. The sample size of Exp. 1 was estimated based on the effect size (Cohen’s d from a t-test) of serial dependence in visual time perception observed in another study from our group (Togoli et al., in preparation). Namely, we computed an average estimate of the expected effect size based on previous data, which turned out to be d = 1.02. Assuming a two-tailed distribution and a power of 95%, the resulting minimum sample size was 15 subjects, which we rounded up to 20 to account for possible excluded participants. In Exp. 2, we based our estimate of effect size on another recent study from our group (Fornaciai et al., 2020) investigating the neural signature of perceptual history across different magnitude dimensions, including time. The neural signature of serial dependence in time perception in this study was explored used a multivariate decoding procedure, which in the case of time perception showed an average effect size (Cohen’s d, again based on a t-test) of 0.8. Based on this effect size, a two-tailed distribution, and a power of 95%, we estimated a minimum sample of 19 subjects, which we conservatively increased to 25 to account for the possible exclusion of a few participants. None of the experiments reported here was pre-registered.

### Apparatus and stimuli

All the experiments were performed in a quiet and dimly lit room. Visual stimuli were presented on a 1920 x 1080 monitor screen running at 120 Hz, positioned at a distance of about 80 cm from the eyes of the participant. The auditory stimuli were presented through loudspeakers positioned behind the screen, at a distance of 50 cm from each other. Visual stimuli were circular noise textures, that could either be composed of black and red squares, or black and green squares (see below *Procedure*). All the stimuli were generated using the Psychophysics Toolbox (Kleiner et al., 2007; Pelli, 1997) on MatLab (version r2015b; The Mathworks, Inc.). The circular area of the visual textures had a radius of 200 pixels (∼3.45 degrees of visual angle, deg, from a viewing distance of 80 cm). The auditory stimuli had an intensity of ∼65 dB measured from the position of the participant. In both Exp. 1 and Exp. 2, visual stimuli were presented either on the left or on the right of a central fixation point, centred on the middle horizontal plane of the screen, and with a horizontal eccentricity (centre to centre) of 12 cm. Auditory stimuli were pure tones, defined by a frequency of either 700 Hz, or 1100 Hz. The tones had a 2-ms ramp at the onset and offset to avoid sound distortions and clicks. In the “feature-selectivity” condition of Exp. 1 and Exp. 2, the tones were presented through both the speakers, while in the “spatial-selectivity” condition the sounds were presented through only one of the speakers, i.e., either the right or the left one.

### Procedure

In Exp. 1, participants performed two different conditions in two different modalities, in a 2 × 2 within-subject design. The two conditions were the “feature-selectivity” condition, where we tested the selectivity of the behavioural signature of perceptual history (i.e., serial dependence effect) for the features of successive stimuli, and a “spatial-selectivity” condition, where we tested the selectivity of the effect for the position of successive stimuli. These two conditions were performed with auditory or visual stimuli in two separate parts of the experiments performed within the same day.

Across all experiments, participants performed a duration discrimination task, comparing the duration of a constant reference stimulus (200 ms) against a variable probe with different durations randomised across trials (100, 140, 200, 280, or 400 ms). To induce perceptual history effects, we presented a task irrelevant “inducer” stimulus before the reference, which could be either 100 ms or 400 ms long, following a similar procedure already used in previous studies (e.g., (Fornaciai & Park, 2018b; Togoli et al., 2021)). Each trial started with participants keeping their gaze on a central fixation point. First, the inducer stimulus was presented on the screen, followed by the reference after an inter-stimulus interval (ISI) of 200-250 ms, and finally the probe after an ISI of 350-450 ms. After the offset of the last stimulus, the fixation point turned red, signalling to the participants that the stimulus presentation was over and to provide a response. Participants were asked to compare the reference and the probe and to report which one between these two stimuli lasted longer in time, by pressing the appropriate key on a keyboard (either “2” or “3,” to indicate whether the second or third stimulus in the sequence was longer). After providing a response, the next trial started automatically after 400-500 ms. When explaining the task, the participants were told that the first stimulus in each sequence (i.e., the inducer) was always irrelevant to the task. In each sub condition of the experiment (i.e., auditory feature-selectivity, auditory spatial-selectivity, visual feature-selectivity, and visual spatial-selectivity), each participant completed 5 blocks of 112 trials, for a total of 56 combinations of the levels of inducer, inducer features/position (i.e., same/different features or same/different position), and probe stimulus duration. All the conditions were performed in a single day. Participants performed a few practice trials (5-10) before the start of the experiment, to ensure that they correctly understood the procedure, and never received feedback about their responses in the main experimental session.

In the feature-selectivity condition, we manipulated whether the features of inducer and reference were similar or different, intermixing “same” and “different” trials within the same blocks. In audition, we chose to modulate the frequency (i.e., pitch) of the stimuli. Namely, while the pitch of reference and probe was always 700 Hz, the pitch of the inducer could be either 700 Hz (same features) or 1100 Hz (different features). In vision, we chose to modulate the colour of the stimuli. In this case, while the reference and probe stimuli were always black and red noise textures, the inducer could either be a black and red texture (same colour) or a black and green texture (different colour). Pitch and colour were chosen to define the features of the stimuli as they are easily and immediately discriminable, and hence there is no uncertainty about whether the stimuli were the same or different. In this condition, all the stimuli were always presented in the same spatial position. Namely, auditory stimuli were presented through both the left and right speaker, while visual stimuli were presented either on the left or on the right of the central fixation point. In the spatial-selectivity condition, we instead manipulated the position of the stimuli and the spatial match of inducer and reference. In this case, all the auditory tones had a pitch of 700 Hz, and all the visual stimuli were black and red textures. In the auditory modality, the reference and probe were presented from either the left or the right speaker, randomised across trials. The inducer was thus presented either from the same speaker as the other stimuli (same position) or from the other speaker (different position). The same procedure was employed in vision. Namely, the refence and probe stimuli could be presented either on the left or on the right of the central fixation point (randomised across trials), and the inducer could be presented either in the same spatial position as the other stimuli (same position), or on the opposite portion of the screen (different position).

In Exp. 2, we used the same paradigm as in Exp. 1, with the addition of electroencephalographic (EEG) recording, but with a different experimental design. Namely, due to the longer time needed to carry out the EEG experiment, the feature-selectivity and spatial-selectivity condition were performed by two independent groups of participants, each including 25 participants. Another difference compared to Exp. 1 was the ISI across the different stimuli, which was adapted to the EEG technique to ensure large enough epochs for the analysis. Namely, the ISI between inducer and reference was 500-600 ms, while the ISI between reference and probe was 800-900 ms. The inter-trial interval was 300-400 ms. In Exp. 2, in each sub-condition, participants performed 5 blocks of 100 trials, for a total of 50 repetitions of each combination of inducer duration, features/position (same or different), and probe duration. Besides these differences, the procedure was identical to Exp. 1.

### Behavioural data analysis

In both Exp. 1 and Exp. 2, we analysed the participants’ performance in the duration discrimination tasks to assess to what extent the perceived duration of the reference stimulus was affected by the preceding inducer, and whether this effect was modulated by the features or the relative position of the stimuli. The proportion of responses in the task (“probe longer”) obtained from each participant in each condition and modality was fitted with a cumulative Gaussian (“psychometric”) function according to the maximum likelihood method (Watson, 1979). In order to improve the psychometric fit, we applied a finger error rate correction (2.5%) to account for random errors and lapses of attention (Wichmann & Hill, 2001). From the psychometric fit, we then computed the measures of accuracy and precision in the task. The “point of subjective equality” (PSE), which represents the perceived duration of the reference stimulus (i.e., accuracy), was defined as the median of the psychometric fit. To assess the precision in the task, we first computed the just noticeable difference (JND) as the difference in probe duration between the 50% and 75% “probe longer” response levels. Moreover, as an additional and more accurate measure of precision, we computed the Weber’s fraction, which represents a measure of precision taking into account the perceived magnitude of the stimuli (WF = JND/PSE). PSE, JND, and WF measures were computed separately for each level of inducer duration and condition. Based on the WF, two participants were excluded from the feature-selectivity condition of Exp. 2 as they showed excessively poor performance in the task (WF > 2).

To better assess and compare the serial dependence effects obtained in the different modalities as a function of the properties of the stimuli, we computed a “serial dependence effect” index based on the normalised difference between PSEs as a function of inducer duration, according to the following formula:

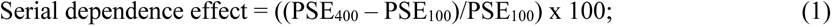

Where PSE_100_ refers to the PSE obtained when the reference was preceded by the shorter inducer (100 ms), while PSE_400_ refers to PSEs obtained with the longer inducer (400 ms). This index was calculated separately for each participant and condition, and the average is shown in Fig. 2. The differences in the serial dependence effect across modalities and conditions were assessed using a linear mixed effect model regression, entering “modality” (vision vs. audition), “condition” (feature vs. spatial selectivity), and “congruency” (same vs. different, either in terms of features or position) as fixed effects, and the subjects as random effect. This regression model was then followed up by a series of paired and one-sample t-tests to better characterise the pattern of effects.

### Electrophysiological recording and analysis

In Exp. 2, the electroencephalogram (EEG) was recorded throughout the experimental session, using the Biosemi ActiveTwo system and a 32-channel cap based on the 10-20 system layout. The EEG signals were recorded with a sampling rate of 2048 Hz. In addition to the 32 electrodes mounted on the cap, we also measured the electro-oculogram (EOG) via a channel attached below the left eye of the participant. The electrode offsets across channels were usually kept below 20 µV, but occasional values up to 30 µV were tolerated.

EEG pre-processing and data analysis was performed offline in Matlab (version R2021b), using the EEGLAB (Delorme & Makeig, 2004) and ERPlab (Lopez-Calderon & Luck, 2014) packages. During pre-processing, EEG recordings were first high-pass filtered (0.1 Hz) and re-referenced to the average of all the channels used. The continuous EEG data were then segmented into epochs spanning from - 200 ms to 800 ms after stimulus onset, time-locking the epochs to the onset of the reference stimulus and sorting them as a function of the duration and properties of the preceding inducer. The pre-stimulus interval (-200:0 ms) was used to perform baseline correction, in order to subtract any spurious effect related to the processing of the inducer stimulus itself from the response evoked by the reference. To clean up the data from noise and artifacts, we performed an independent component analysis (ICA). Furthermore, we applied a step-like artifact rejection procedure (amplitude threshold = 40 μV, window = 400 ms, step = 20 ms) aimed to further remove any remaining large artifact from the EEG signals, which yielded an average (± SD) rejection rate of 1% ± 1.87% of the trials. Finally, we applied a lowpass filter with a cut-off of 30 Hz before computing the ERPs.

To assess the neural signature of perceptual history, we performed a series of analyses on ERP and EEG data. First, as a measure of the effect of perceptual history, we computed the difference (or contrast) in ERP amplitude as a function of the duration of the inducer, irrespective of the properties of the stimuli, throughout all the EEG channels. The scalp topography of this contrast amplitude was then used to select the channels of interest used for further analysis. To improve the signal-to-noise ratio, we selected two channels and used their average for further analyses (i.e., the channel showing peak contrast amplitude and an adjacent channel showing the highest amplitude among the surrounding electrodes). In the spatial-selectivity condition, we similarly selected the best channel and a nearby one, but this time considering the scalp locations contralateral to the position of the reference stimulus (i.e., two channels in the left hemisphere when the reference was presented on the right, and vice versa). In the feature selectivity condition, we selected T8 and P8 in audition, and P8 and PO4 in vision. In the spatial-selectivity condition, we selected T8 and P8 in the right hemisphere and T7 and P7 in the left hemisphere, in both the auditory and visual condition.

In our main ERP analysis, we assessed the contrast amplitude – which was considered as a neural signature of the perceptual history effect induced by the inducer stimulus – separately for the different features or positions of the stimuli. First, to further increase the signal-to-noise ratio, we averaged the contrast amplitude across a series of 100-ms time windows, with a step of 25 ms. To compare the contrast amplitude across conditions where the inducer had either the same or different features or position, we performed a series of paired t-tests. To control for multiple comparisons, we used a cluster-based non-parametric test, which is described in details below. Then, to assess the extent to which the contrast amplitude relates to the serial dependence bias measured behaviourally, we performed a linear mixed-effect regression analysis. In this analysis, we entered the difference in PSE corresponding to the 400-ms and 100-ms inducer (ΔPSE) as dependent variable. As fixed effect factors, we entered the contrast amplitude (ΔERP), the match between inducer and reference (same vs. different, either features or position), and the interaction between these two factors. The subjects were added to the analysis as the random effect. This analysis was performed independently across each time window throughout the reference epoch. Again, multiple comparisons were controlled by performing a cluster-based non-parametric test.

Moreover, we also performed a non-linear regression analysis, considering the amplitude of EEG responses across individual trials rather than the averaged ERPs. In this analysis, we thus entered the trial-by-trial EEG amplitude as dependent variable, and assessed its relationship with the inducer and its properties, and the response that participants provided in each trial. In the feature-selectivity condition, we entered as factors the duration of the inducer (100 vs. 400 ms), the features of the inducer (same vs. different compared to the reference), and the behavioural response, i.e., the binary choice (“reference longer” or “probe longer,” coded as 0 and 1) made by participants at the end of each trial. In the spatial-selectivity condition, we entered again the duration of the inducer and the behavioural response as factors, with the addition of the reference position (i.e., reference presented on the left or on the right of the fixation point), and the spatial match of the inducer (same position vs. different position). This non-linear analysis was used to assess the contribution of each of those factors to the amplitude of the trial-by-trial EEG response, which was characterised in terms of beta value.

The beta value indicates the extent to which the EEG response changes (in terms of μV of amplitude) across the different levels of each factor, and hence it could provide evidence for a relationship between the EEG activity, the experimental manipulations, and the participants’ response in each trial. Similarly to the linear regression analysis, the non-linear regression was performed independently at each time window across the reference epoch. Beta values across the group, at each time window were then assessed using one-sample t-tests against the null hypothesis of zero effect. This series of tests was again controlled using a cluster-based non-parametric test.

Across the different tests performed during the ERP/EEG analysis, we controlled for multiple comparisons using a cluster size threshold and a cluster-based non-parametric tests. First, to be considered as a potentially significant cluster, we set a threshold of at least three consecutive significant time windows from the original test performed (i.e., three consecutive significant paired t-tests, for example). In the non-parametric test, each cluster of consecutive time windows was compared to the clusters that could be obtained across several repetitions of the same test but with either randomised data or by performing permutations of the data being compared. Regarding the cluster-based procedure in the case of the paired t-tests, we split the data corresponding to the two conditions being compared into two randomly selected subsets, and created two new sets of data by concatenating two halves corresponding to the two original conditions. We then performed the paired t-test across these intermixed datasets. This procedure was performed across each time window within each of the clusters observed in the actual analysis, and repeated 10,000 times. To test for the significance of the actual cluster, we set a cluster-level threshold equal to the minimum t-value observed in the actual cluster, and assessed how many consecutive time windows exceeded this threshold within the simulated cluster. We then counted how many times the simulation yielded a cluster of consecutive time windows equal to the actual cluster, and defined the cluster-level p-value as the proportion of times that the same number of consecutive time windows was observed across the 10,000 simulations. In the case of the linear regression analysis, the procedure was identical, but to compute the simulated clusters we randomly shuffled the design matrix containing the values of the predictors. Finally, in the case of the one-sample t-tests used to assess the significance of beta values from the non-linear analysis, we flipped the sign of a random half of data before performing the one-sample t-test.

### Data availability

All the data generated during the experiments described in this manuscript is freely available on Open Science Framework at this link: https://osf.io/7ex9y/ (DOI: 10.17605/OSF.IO/7EX9Y).

## ACKNOWLEDGEMENTS

This project has received funding from the European Union’s Horizon 2020 research and innovation programme under the Marie Sklodowska-Curie grant agreement No. 838823 “NeSt” to MF, from the European Research Council (ERC) under the European Union’s Horizon 2020 research and innovation programme grant agreement No. 682117 BIT-ERC-2015-CoG to DB, and from the Italian Ministry of University and Research under the call FARE (project ID: R16X32NALR) and under the call PRIN2017 (project ID: XBJN4F) to DB.

